# Sex-specific proteomic analysis of epileptic brain tissues from Pten knockout mice and human refractory epilepsy

**DOI:** 10.1101/2025.03.27.645753

**Authors:** Yibo Li, Zahra Sadri, Katherine J. Blandin, David A. Narvaiz, Uma K. Aryal, Joaquin N. Lugo, Nicholas P. Poolos, Amy L. Brewster

## Abstract

**Rationale:** Epilepsy presents significant sex-based disparities in prevalence and manifestation. Epidemiological studies reveal that epilepsy is more prevalent in males, with lesional types being more common, whereas idiopathic generalized epilepsies are more frequently observed in females. These differences stress the importance of considering sex-specific factors in epilepsy diagnosis, treatment, and mechanistic research using preclinical models. To elucidate potential molecular differences that could explain these disparities and inform personalized treatment strategies, we conducted a proteomic analysis of epileptic brain tissues from both an experimental mouse model of genetic epilepsy and humans with drug-resistant epilepsy (DRE).

**Methods:** We employed mass spectrometry-based proteomic analysis on brain tissues from DRE patients and the *Pten* knockout (KO) mouse model of genetic epilepsy with focal cortical dysplasia. Mouse samples included hippocampi from adult wild-type (WT) and *Pten* KO mice (4-5 per group and sex). Human samples included temporal cortex from 12 DRE adult patients (7 males, 5 females) and 5 non-epileptic (NE) controls (2 males, 3 females). Brain biopsies were collected with patients’ informed consent under approved IRB protocols (Indiana University Health Biorepository). Proteomic profiles were analyzed using principal component analysis (PCA) along with volcano plots to identify significant changes in protein expression. The enrichment analysis of differentially expressed proteins was conducted by Gene Ontology (GO) and Kyoto Encyclopedia of Gene and Genomes (KEGG) pathway.

**Results:** PCA revealed distinct clustering of brain proteomes between epilepsy and control cases in both human and mice, with 390 proteins showing significant differences in human and 437 proteins in mouse samples. These proteins are primarily associated with ion channels, synaptic processes, and neuronal energy regulation. In the mouse model, males have more pronounced proteomic changes than females, with enrichment in metabolic pathways and VEGF signaling pathway, indicating a more severe vascular permeability impairment in males. In human DRE cases, 118 proteins were significantly changed by comparing epileptic females to males. Pathway analysis revealed changes in metabolic pathways and the HIF-1 signaling pathway, indicating that altered neuronal activity and inflammation may lead to increased oxygen consumption.

**Conclusion:** These findings highlight significant differences between epilepsy and control brain samples in both humans and mice. Sex-specific analysis revealed distinct pathway enrichments between females and males, with males exhibiting a broader range of alterations, suggesting more extensive proteomic alterations. This study offers valuable insights into potential underlying mechanisms of epilepsy and underscores the importance of considering sex as a key factor in epilepsy research and therapeutic development.

## Introduction

Epilepsy is a heterogeneous seizure disorder characterized by the occurrence of spontaneous seizures that can coexist with other neurologic problems (1). Epilepsy impacts ∼65 million people worldwide and 3.4 million individuals in the United States (2). One-third of people with epilepsy experience drug-resistant seizures (3), and up to 80% of those with drug-resistant epilepsy (DRE) suffer from cognitive impairments (4, 5). Despite being one of the oldest recognized disorders (6), the exact causes remain unknown in more than half of the cases. Moreover, identifying and understanding the causal mechanisms of epilepsy is complicated by differences between males and females with epilepsy (7). Epidemiological studies indicate sex-based differences in epilepsy prevalence, presentation, and treatment response (8–11). While both males and females have predominantly focal epilepsies, males exhibit higher overall epilepsy rates, an increased seizure burden, and a slightly higher propensity toward epilepsy associated with injuries (7–9, 12–15). In contrast, females more often have idiopathic generalized epilepsies, experience seizures preceded by auras, and are more susceptible to adverse reactions from anti-seizure medications (7–9, 12–15). This sex-based disparity stresses the importance of investigating the underlying molecular causes.

Despite these well-documented sex differences in human epilepsy (7), discovery-based and translational research utilizing omics approaches to pinpoint potential underlying mechanisms has largely focused on male subjects or mixed-sex cohorts (for review see (16)), with comparatively fewer studies explicitly analyzing sex-specific molecular differences (17). The male-biased research paradigm has created a critical knowledge gap (7, 13), limiting our ability to develop efficacious therapeutic strategies that take into consideration sex-specific molecular profiles. Emerging evidence from rodent models of genetic and acquired epilepsies suggests that males and females exhibit distinct pathophysiological phenotypes (7, 17–22). Recent studies from genetic models of epilepsy have shown that female epileptic mice have higher seizure resilience and increased survival compared to male epileptic mice (18–20, 23). Furthermore, only one study has shown a comprehensive characterization of male and female mice using transcriptomic analysis in genetically epileptic mice, where distinct pathway alterations were found in males and females, with pathways related to metabolism enhanced in females and broader changes in pathways affecting synaptic and glial structure and function in males (18). These distinct molecular signatures suggest that males and females may utilize different mechanisms to develop and maintain epileptic networks.

To gain more knowledge on the molecular profiles of males and females with epilepsy, in the current study, we examined the hippocampal proteome in the *Pten* knock out (KO) mouse model of genetic epilepsy, which recapitulates the human epilepsy associated with focal cortical dysplasia (FCD) (24). *Pten* KO mice are characterized by enhanced mTOR activation, enlarged dentate gyrus cortical, and develop epilepsy by postnatal week 2 (24–30), making them a valuable model for investigating molecular pathways underlying epilepsy. Given the limited number of studies directly comparing male and female molecular signatures in epilepsy, we also analyzed differentially expressed proteins in human epileptic and non-epileptic brain samples, with a specific focus on sex differences in protein expression. Our findings aim to provide insights into sex-specific molecular alterations in epilepsy, which may facilitate the development of more precise and effective therapeutic interventions tailored to male and female patients.

## Materials and Methods

### Ethics statement

Human brain samples were collected with patients’ informed consent under approved Institutional Review Board (IRB) protocol #1011004282 (Development of a Biorepository for Methodist Research Institute; Indiana University Health Biorepository). All identifiable information was removed before conducting experiments and analyses under IRB protocol # 1507016240 (Purdue University) and IRB protocol # 21-126 (Southern Methodist University). Following collection, human brain samples were stored at -80°C.

### Human brain tissue

Cortical tissue samples were obtained from two groups: patients with drug-resistant epilepsy (DRE) who underwent resective surgery (n = 12; 5 females, 7 males) and non-epileptic (NE) patients undergoing tumor resection (n = 5; 3 females, 2 males). The NE samples were collected from cortical tissue adjacent to tumors, including glioblastoma, and were confirmed as relatively normal by neuropathological examination. This tissue, routinely removed to access deep-seated tumors, was obtained from patients with no history of seizures. All samples were from temporal lobe cortex, with no hippocampal tissue included. Age ranges were comparable across groups: DRE females (21-63 years, mean 37.2 ± 15.26), DRE males (20-68 years, mean 41.6 ± 18.05), NE females (53-69 years), and NE males (28-51 years). The mean duration of epilepsy in DRE patients was 16.12 ± 10.3 years for females and 24.13 ± 18.89 years for males.

**Table 1.**
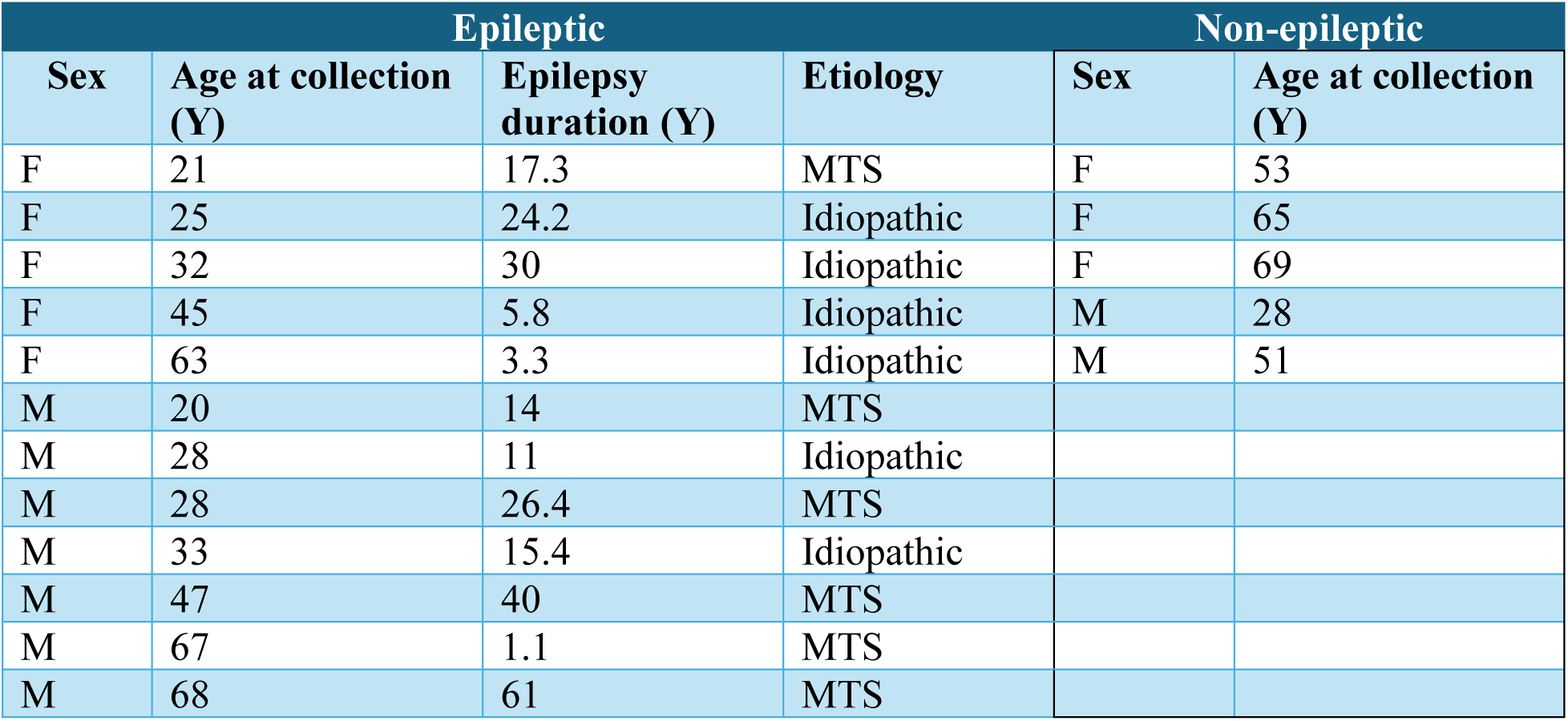
Clinical characteristics of study subjects, including sex, age at sample collection (years, Y), epilepsy duration (years, Y) and etiology are shown for the epileptic (E) group. Demographic information (sex and age) is shown for the non-epileptic (NE) group. Abbreviation: F, female; M, male; MTS, mesial temporal sclerosis.

### Mouse samples

Neuronal subset-specific wildtype (WT) and *Pten* knockout (KO) mice (RRID: MGI:371406)(2–4) (2-3 months old) (25–27) of both sexes were used for this study (n = 4–5 per group). Mice were euthanized with a lethal dose of Beuthenasia (200 mg/kg i.p.), and hippocampi were collected, frozen in dry ice, and stored at -80°C until used. Mice were housed in cages under standard environmental conditions (12h light-dark cycle, temperature 22°C, free access to food and water) and bred according to Baylor University’s Institutional Animal Care and Use Committee and the National Institute of Health’s *Guide for the Care and Use of Laboratory Animals* (Protocol # 971364–9). Our research followed NIH, Institutional, and ARRIVE guidelines (31).

### Proteomics sample preparation and data analysis

Brain tissues were homogenized in 1X PBS and the whole tissue homogenates were submitted to the Proteomics core facility. Samples were then placed in 100 mM ABC and homogenized in precellys homogenization vials (Bertin Technologies SAS, France) for 90 seconds at 6500 rpm. Protein concentration was then measured by bicinchoninic acid (BCA) assay (Pierce Chemical Co., USA) and 50 ug protein (equivalent volume of tissue lysates) were precipitated at -20 °C overnight with 4 volumes of cold (-20 °C) acetone. Samples were then centrifuged at 13,500 × *g* for 15 minutes, acetone supernatant was discarded, and samples were briefly dried in a vacuum centrifuge before resuspending in 10 uL of 10 mM dithiothreitol (DTT) in 8M urea solution and incubated at 37 °C for one hour for reduction. An equal volume of alkylating mixture (195 uL acetonitrile, 1 uL triethylphosphate (TEP), 4 uL Iodoethanol) was added, and samples were incubated at 37 °C for one additional hour, and consecutively dried in a vacuum centrifuge. Samples were then digested with Lys-C/Trypsin in 50 mM ammonium bicarbonate (ABC) in a 1:25 enzyme to protein ratio using a barocycler (60 cycles: 20 kPSI for 50 seconds and 1 ATM 10 seconds, at 50 °C). Digested peptides were desalted using C_18_ Silica MicroSpin Columns (The Nest Group, Inc. USA), eluted cleaned peptides were dried in SpeedVac and resuspended in 3% acetonitrile/96.9% MilliQ, and 0.1% formic acid (FA).

### Liquid Chromatography-Tandem Mass Spectrometry (LC-MS/MS) analysis

Peptides were analyzed in a Dionex UltiMate 3000 RSLC nano System (Thermo Fisher Scientific, Odense, Denmark) coupled with Orbitrap Fusion Lumos Tribrid Mass Spectrometer (Thermo Fisher Scientific, Waltham, MA, USA) as described previously (32). Briefly, reverse phase peptide separation was accomplished using a trap column (300 μm ID × 5 mm) packed with 5 μm 100 Å PepMap C18 medium coupled to a 50-cm long × 75 µm inner diameter analytical column packed with 2 µm 100 Å PepMap C18 silica (Thermo Fisher Scientific) with the column temperature maintained at 50°C. Mobile phase solvent A was 2% acetonitrile (ACN), 98% water and 0.1% Formic Acid (FA). Mobile phase solvent B was 80% ACN, 20% water and 0.1% FA. Sample was loaded to the trap column in a loading buffer (3% acetonitrile, 0.1% FA) at a flow rate of 5 uL/min for 5 min and eluted from the analytical column at a flow rate of 200 nL/min using a 160-min LC gradient as: 6.5 to 27% of solvent B in 82 min, 27-40% of B in next 8 min, 40-100% of B in 7 min at which point the gradient was held at 100% of B for 7 min before reverting back to 2% of B, and hold at 2% of B for next 15 min for column equilibration. The column was further washed and equilibrated by using three 30-min LC gradient before injecting the next sample. All data were acquired in the Orbitrap mass analyzer and data were collected using an HCD fragmentation scheme. For MS scans, the scan range was from 350 to 1600 *m*/*z* at a resolution of 120,000, the automatic gain control (AGC) target was set at 4 × 10^5^, maximum injection time 50 ms, dynamic exclusion 30s and intensity threshold 5.0 ×10^4^. MS data were acquired in Data Dependent mode with cycle time of 5s/scan. MS/MS data were collected at a resolution of 15,000.

### Data Analysis

LC-MS/MS data were analyzed using MaxQuant software (version 1.6.3.3) against the combined non-redundant protein sequence Uniprot database for human and mouse (www.uniprot.org) for protein identification and label-free relative quantitation. The following parameters were used for database searches: precursor mass tolerance of 10 ppm; enzyme specificity of trypsin/Lys-C enzyme allowing up to 2 missed cleavages; oxidation of methionine (M) as a variable modification and iodoethanol (C) as a fixed modification. The false discovery rate (FDR) of peptide spectral match (PSM) and protein identification were set to 0.01. Proteins with LFQ # 0 and MS/MS (spectral counts) ≥2 were only considered true identification and used for downstream statistical analysis and data visualization.

### Proteomics computational analysis

For the subsequent statistical analyses, intensity-based absolute quantitation (iBAQ) for both human and mouse samples was analyzed, and the dataset was limited to proteins quantified in at least half of the samples in each condition. The missing values were imputed by using K-nearest neighbors (KNN) and Z-score normalization was performed. Volcano plot, principal components analysis (PCA), and hierarchical cluster analysis were performed using R software (version 4.4.0) following similar studies (33).

### Bioinformatics analysis

For hierarchical clustering, Euclidean distance was used as the distance measure, and complete linkage was used to combine clusters. The ShinyGO 0.76, which is a web-based tool (http://bioinformatics.sdstate.edu/go/), was used to conduct enrichment analysis of differentially expressed proteins (DEPs) by which Gene Ontology (GO) and Kyoto Encyclopedia of Gene and Genomes (KEGG) pathway were analyzed. The GO classification consisted of biological processes (BP), molecular functions (MF), and cellular components (CC). *P* value < 0.05 was set as the threshold of significance. For KEGG pathway enrichment analysis, the information of DEPs was mapped into the KEGG database to obtain the enriched pathway.

### Statistical analysis

The differences between the two groups were evaluated using the Student’s *t*-test, the plots were generated in R (version 4.4.0) and GraphPad Prism software (version 10.2.0). Differences were considered statistically significant at *p* < 0.05. Significantly altered proteins are included in the Supplementary Excel file.

### Data availability

Raw data from this study can be accessed from the Texas Data Repository (Accession No. ZMP8KU).

## Results

Proteomics offers a powerful approach to uncovering the complex molecular changes associated with epilepsy by enabling the identification and quantification of differentially expressed proteins in brain tissues. Building upon emerging evidence highlighting sex-specific transcriptomic differences between epileptic and control groups in a genetic mouse model (18), we aimed to explore the potential mechanisms that may be directly linked to epilepsy at the proteomic level. Proteomic profiling was done in hippocampi dissected from *Pten* KO mice, which is a mouse model of genetic epilepsy resembling focal cortical dysplasia (FCD) (24, 34). These were compared to those of WT mice. The PCA analysis of the proteome showed distinct clusters between the WT and *Pten* KO groups (Fig. 2A) (p < 0.0039), suggesting that the overall proteomic profiles differed in brains of epileptic and non-epileptic mice. The volcano plot identified 437 proteins with statistically significant differential abundance, including 192 proteins with increased expression and 245 proteins with decreased expression in the hippocampi of *Pten* KO mice compared to WT mice (Fig. 2B, supplementary Excel file). The heatmap illustrated distinct proteomic profiles between WT and *Pten* KO mice, with proteins exhibiting consistent upregulation or downregulation in one group relative to the other (Fig. 2C). GO analysis of the differentially abundant proteins revealed significant enrichment in biological processes related to synaptic transmission and signaling (Fig. 2D). KEGG pathway analysis indicated enrichment pathways such as the TCA cycle, oxidative phosphorylation, glutamatergic synapses, and other pathways implicated in epilepsy (Fig. 2E), suggesting disruptions in neuronal energy metabolism and synaptic function as key contributors to epilepsy pathophysiology.

**Figure 1.**
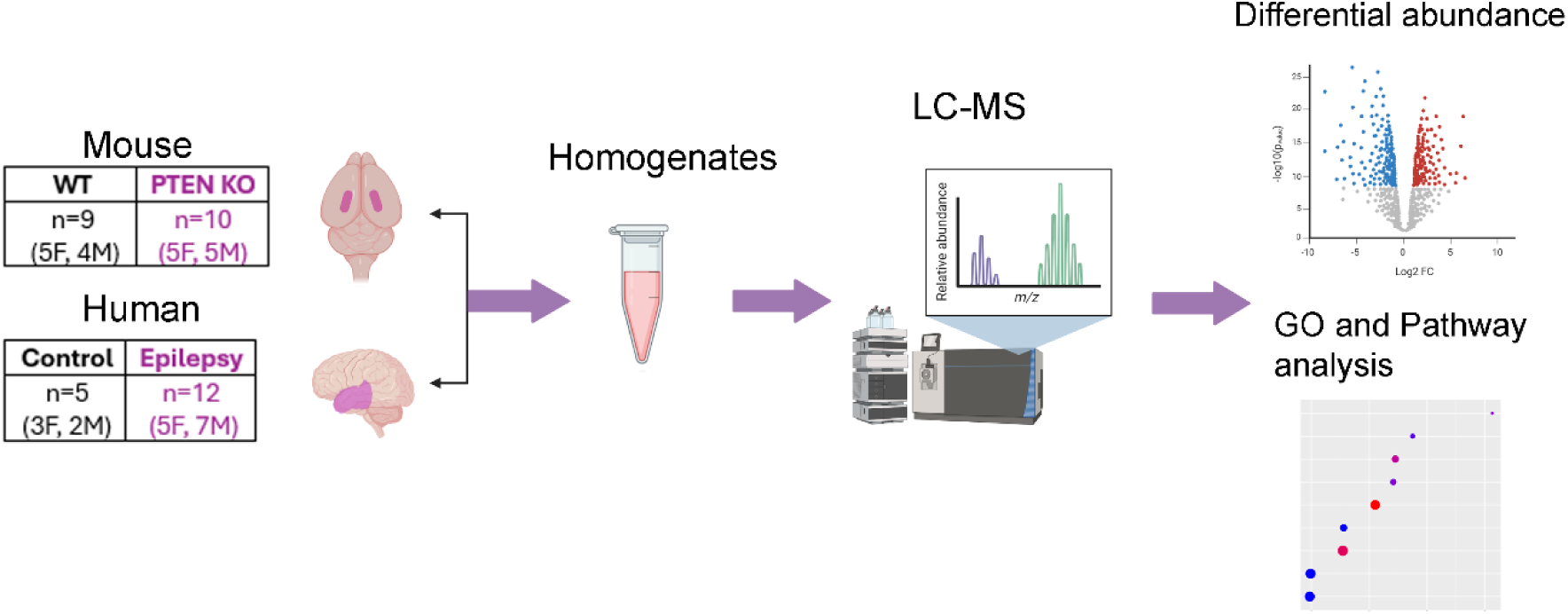
Diagram of the experimental workflow. Mouse and human brain tissues were collected from control and epilepsy groups (Mouse: control, n = 9; epilepsy, n = 10; Human: control, n = 5; epilepsy, n = 12). Proteins were quantified using liquid chromatography-tandem mass spectrometry (LC-MS), followed by statistical analysis, functional enrichment, and pathway analysis to identify differentially expressed proteins and enriched pathways.

**Figure 2.**
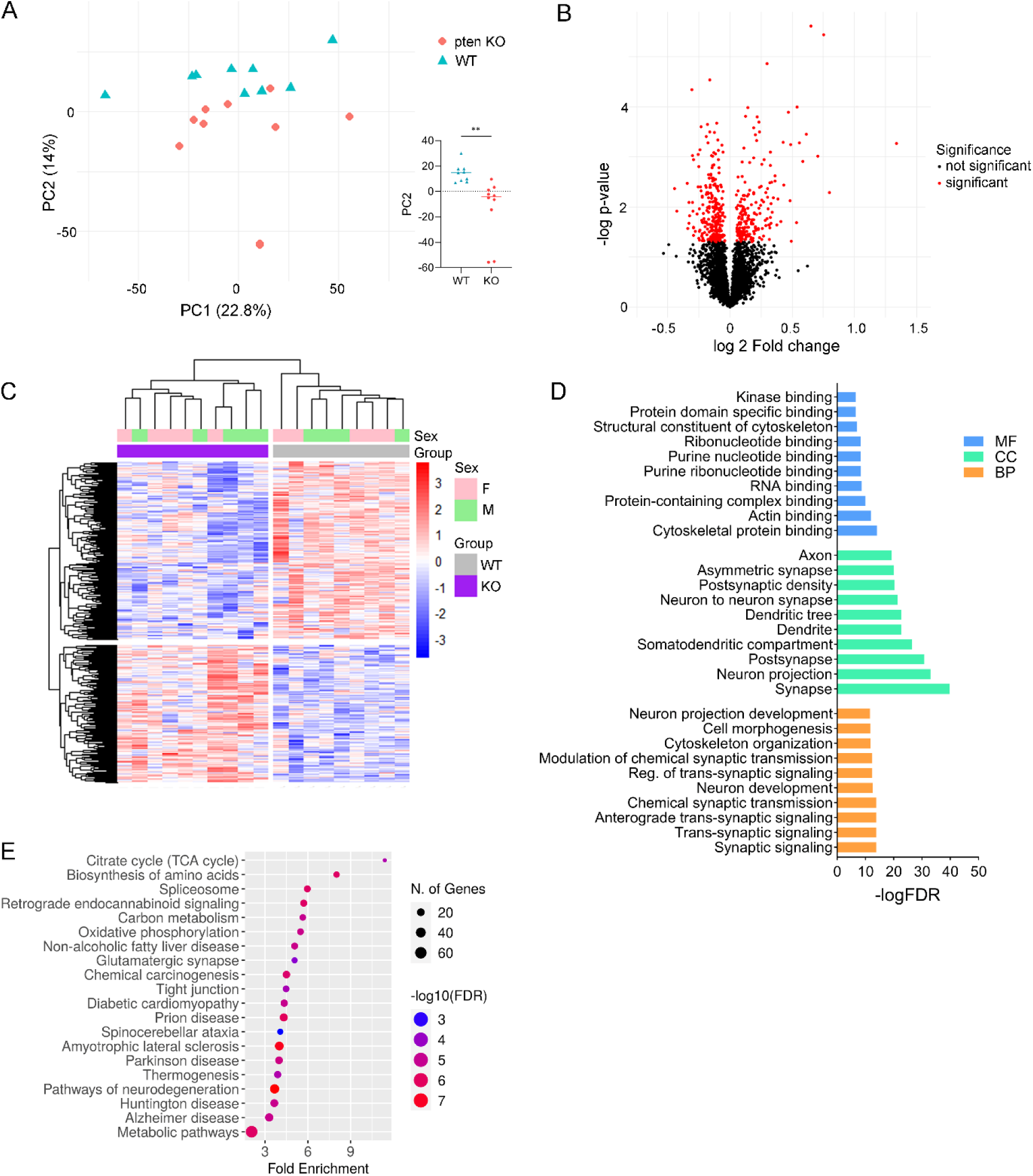
Proteomic differences in the hippocampus of Pten knockout (KO) and wildtype (WT) mice. (A) Principal component analysis (PCA) of Pten KO (red) and WT (blue) groups, showing distinct clustering based on differentially expressed proteins. Each data point represents an individual case. The PC2 bar plot indicates significant separation between groups. (B) Volcano plot displaying protein expression differences for all identified proteins. Proteins highlighted in red showed significantly altered expression in Pten KO mice compared to WT mice. Significance was determined using Student’s t-test. P-values for significantly altered proteins are provided in the Supplementary Table. (C) Heatmap of scaled protein expression values (z-scores), color-coded according to the legend on the right, for significantly changed proteins used in clustering analysis. (D) Gene Ontology (GO) analysis of significantly altered proteins, including biological process (BP), cellular component (CC), and molecular function (MF). The top 10 terms are shown. (E) KEGG pathway analysis of significantly altered proteins. The top 20 pathways are shown.

Evidence suggests that sex can influence disease pathophysiology (7, 11, 17, 18, 35). To investigate whether sex differences contribute to the altered proteome profiles observed in epilepsy, we compared WT-F and WT-M to assess baseline sex differences. PCA showed no statistically significant differences in proteomic profiles (Supplementary Fig. 1A-B). Among 55 significantly altered proteins, 30 were upregulated and 25 downregulated in WT-F (Supplementary Fig. 1C-D), with no KEGG-enriched pathways. Venn analysis revealed minimal overlap between F-WT vs M-WT and F-WT vs F-KO (10 proteins, 2.6%) or M-WT vs M-KO (5 proteins, 1.3%) (Supplementary Fig. 1H), suggesting some baseline sex differences in WT mice.

Next, we compared *Pten* KO mice to their sex-matched WT controls. In females, we identified 168 significantly altered proteins, with 75 upregulated and 93 downregulated in *Pten* KO mice compared to female WT controls (Fig. 3A, 6C). In males, 214 proteins were significantly altered, with 89 upregulated and 125 downregulated in *Pten* KO mice relative to male WT controls (Fig. 3B, 3D). KEGG pathway analysis revealed significant enrichment in protein processing in the endoplasmic reticulum (ER) in females (Fig. 3E). In contrast, males showed significant enrichment in the vascular endothelial growth factor (VEGF) signaling pathway, glutamatergic synapses, Rap1 signaling pathway, among other pathways (Fig. 3F). This suggests a more complex proteomic alteration in males and indicates potential increased vascular permeability, as inferred from VEGF pathway enrichment. Additionally, 14 upregulated proteins were shared between males and females, including S100a6, GFAP, CD44, and VGF (Fig. 3G, 3I). Similarly, 21 downregulated proteins were common to both sexes, such as Syn1 and Palm (Fig. 3H, 3I), indicating that certain molecular processes are consistently dysregulated across the sexes.

**Figure 3.**
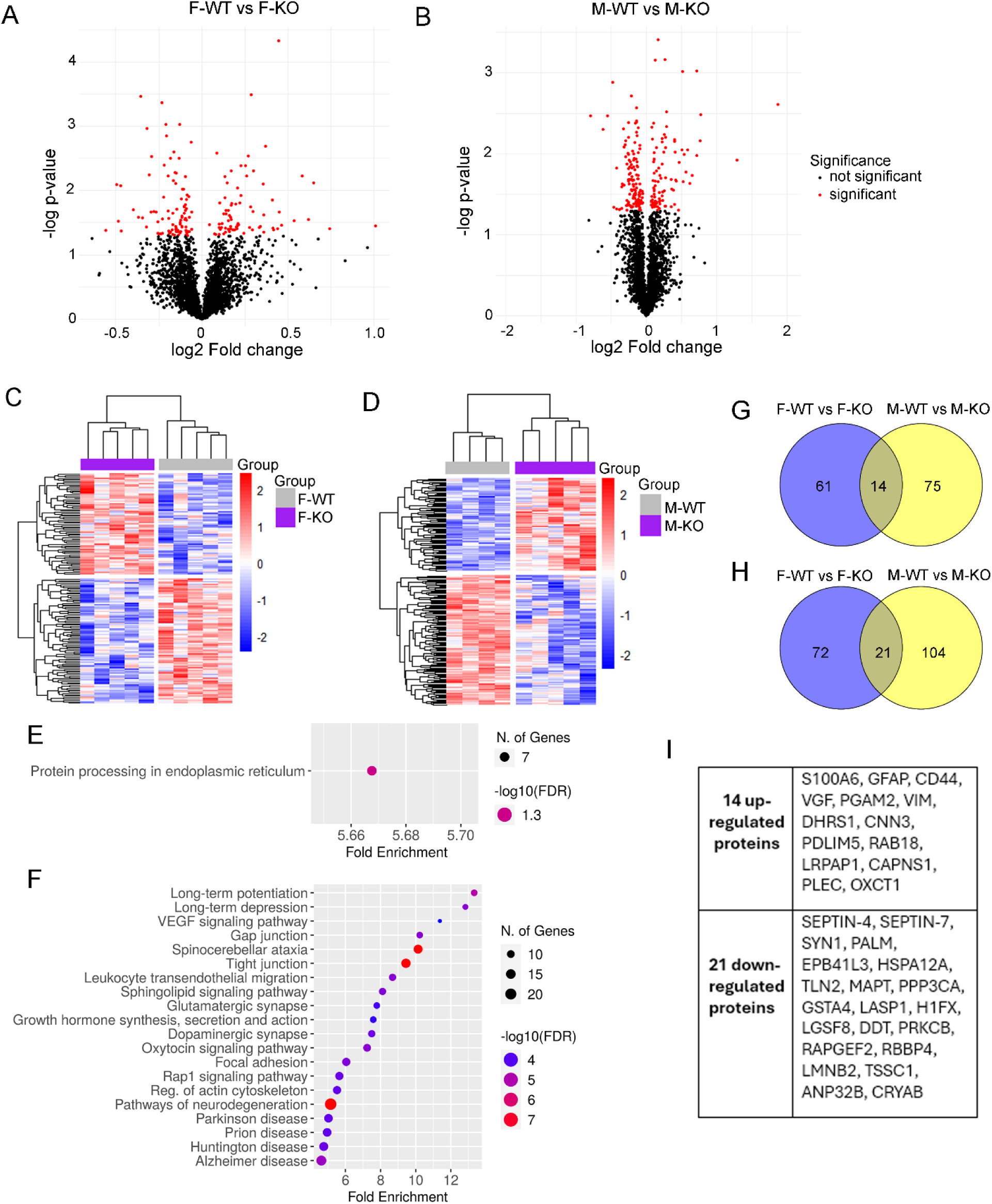
Sex-specific proteomic differences in Pten knockout (KO) and wildtype (WT) mice. (A–B) Volcano plots showing protein expression differences in all identified proteins. Proteins highlighted in red were significantly altered when comparing female KO (F-KO) with female WT (F-WT) (A), and male KO (M-KO) with male WT (M-WT) (B). Significance was determined using Student’s t-test. P-values for significantly altered proteins are provided in the Supplementary Table. (C–D) Heatmaps displaying scaled protein expression values (z-scores), color-coded according to the legend on the right, for significantly changed proteins in F-KO vs F-WT (C) and M-KO vs M-WT (D). (E–F) KEGG pathway analysis of significantly altered proteins in females (E) and males (F). (G) Fourteen upregulated proteins shared between females and males. (H) Twenty-one downregulated proteins shared between females and males. (I) Summary of overlapping proteins (14 upregulated shown in G and 21 downregulated shown in H).

We extended our analysis to human samples to explore the relevance and potential implications of the findings from the *Pten* KO epilepsy model. Consistent with the mouse data, PCA analysis revealed a clear separation between epileptic (E) and non-epileptic (NE) samples (Fig. 4A), indicating distinct proteomic profiles between epilepsy cases and controls in humans. Differential expression analysis identified 390 proteins with significant changes (p < 0.05) between E and NE samples, with 209 proteins upregulated and 181 downregulated in epilepsy cases (Fig. 4B). A heatmap further highlighted distinct expression patterns between E and NE samples (Fig. 4C). Notably, SNCA and VGF were upregulated in epileptic samples (Fig. 4D), consistent with previous reports (36, 37). Gene ontology (GO) analysis of the significantly altered proteins revealed significant enrichment in synaptic transmission and signaling in humans (Fig. 4E). Pathway analysis using KEGG indicated a potential dysregulation of the complement and coagulation cascades, as well as TCA cycle, metabolic dysfunction and synaptic abnormalities in epilepsy (Fig. 4F).

**Figure 4.**
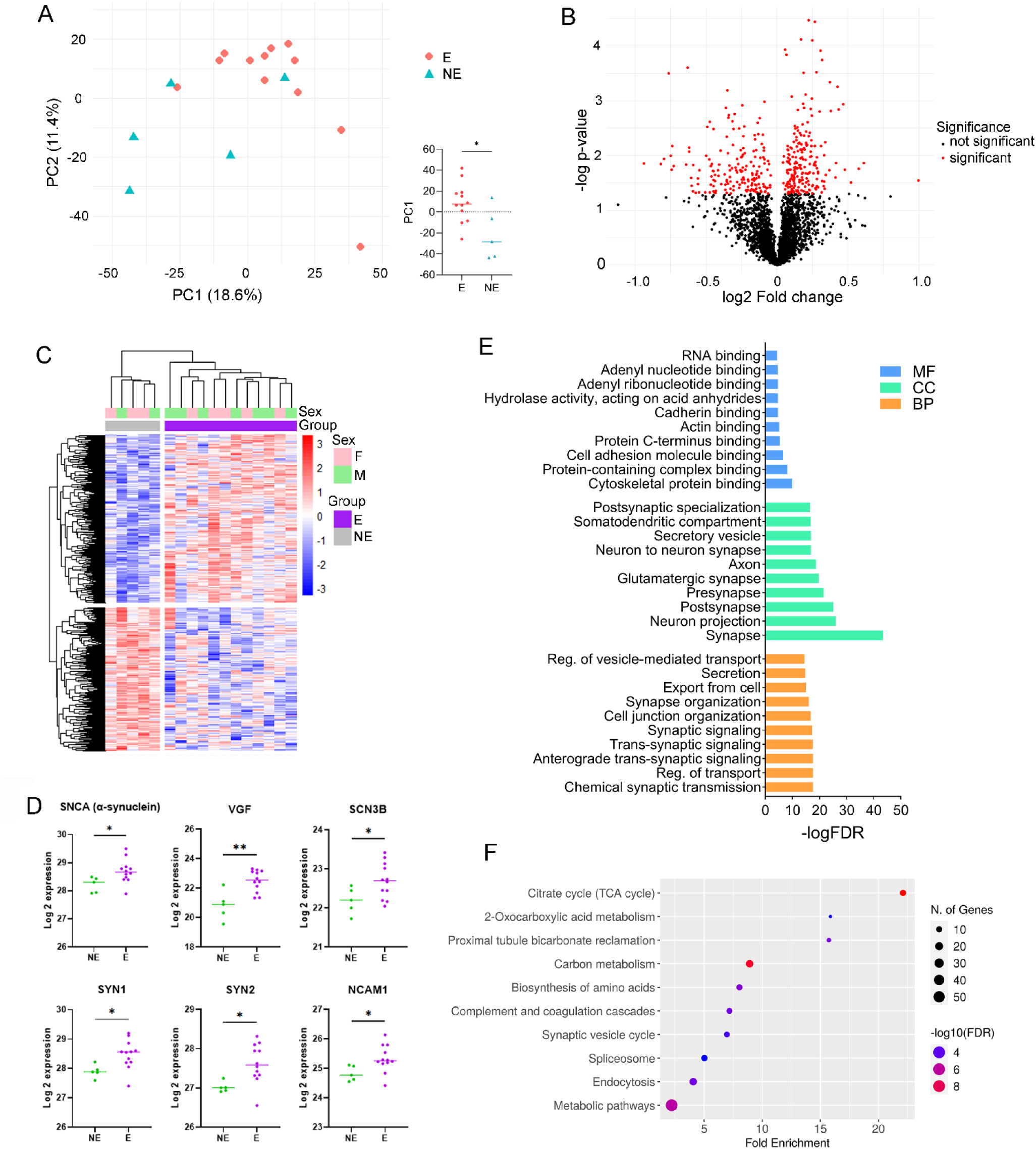
**Proteomic differences between human epilepsy and non-epileptic cases. (**A) Principal component analysis (PCA) of epileptic (E, red) and non-epileptic (NE, blue) cases, showing distinct clustering based on protein expression profiles. Each point represents an individual case. The PC1 bar plot indicates significant separation between groups. (B) Volcano plot displaying protein expression differences for all identified proteins. Proteins highlighted in red showed significantly altered expression when comparing E and NE cases. Significance was determined using Student’s t-test. P-values for significantly altered proteins are provided in the Supplementary Table. (C) Heatmap showing scaled protein expression values (z-scores), color-coded according to the legend on the right, for significantly changed proteins used in clustering analysis. (D) Expression levels of selected proteins of interest with significant alterations in epilepsy. Each point represents an individual case. *p < 0.05; **p < 0.01. (E) Gene Ontology (GO) analysis of significantly altered proteins, including biological process (BP), cellular component (CC), and molecular function (MF). The top 10 terms are shown.(F) KEGG pathway analysis of significantly altered proteins. The top 10 pathways are shown.

To investigate potential sex dimorphism in human epilepsy, we compared the proteomic profiles of males and females with DRE. PCA analysis did not reveal distinct clustering between the two groups (Fig. 5A-B), suggesting that global proteomic profiles are largely similar under the conditions studied. However, differential expression analysis identified 118 proteins with significant changes between females and males in DRE patients, with 69 proteins upregulated and 49 proteins downregulated in females compared to males (Fig. 5C-D). Notably, GFAP and C1qb were differentially expressed between the two groups (Fig. 5E), indicating that while overall proteomic patterns may be comparable, specific proteins exhibit sex-specific regulation in epilepsy. KEGG pathway analysis revealed significant enrichment in several critical pathways, including the HIF-1 signaling pathway, metabolic pathways, and multiple neurodegeneration-related pathways (Fig. 5F). Despite these findings, a key limitation of this study is the limited availability of sex-matched control tissue from individuals without epilepsy. The inclusion of such controls would provide a more comprehensive understanding of epilepsy-specific protein alterations.

**Figure 5.**
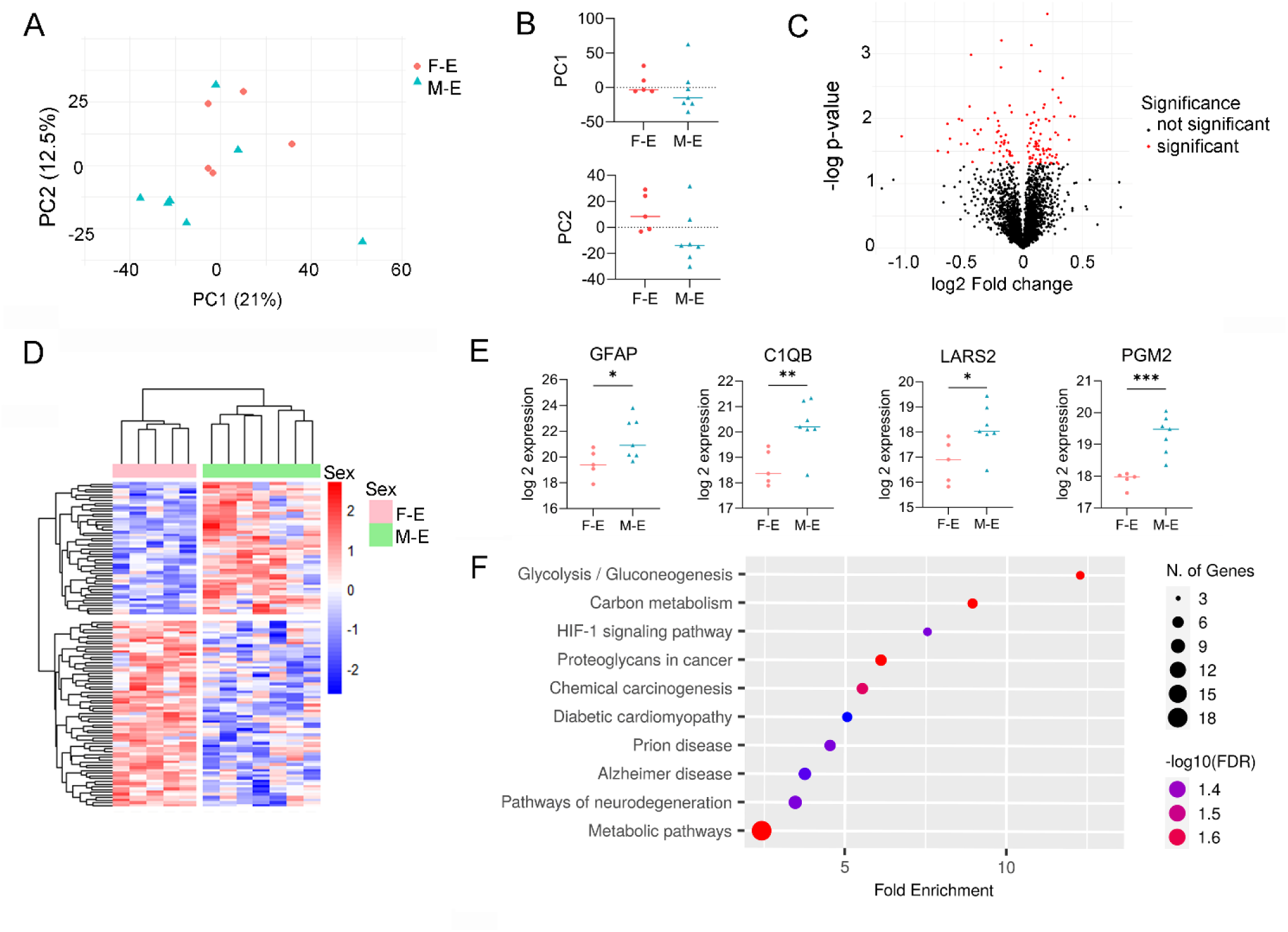
Sex-specific proteomic differences in human epilepsy. (A) Principal component analysis (PCA) of the overall dataset. PCA of epileptic female (F-E, red) and epileptic male (M-E, blue) cases shows similar protein expression profiles. Each point represents an individual case. (B) PC1 and PC2 bar plots indicate no significant separation between F-E and M-E cases. (C) Volcano plot displaying protein expression differences for all identified proteins. Proteins highlighted in red showed significantly altered expression when comparing F-E and M-E cases. Significance was determined using Student’s t-test. P-values for significantly altered proteins are provided in the Supplementary Tables. (D) Heatmap of scaled protein expression values (z-scores), color-coded according to the legend on the right, for significantly changed proteins used in clustering analysis. (E) Expression levels of selected proteins of interest with significant sex-specific differences. Each point represents an individual case. *p < 0.05; **p < 0.01; ***p < 0.001. (F) KEGG pathway analysis of significantly altered proteins. The top 10 pathways are shown.

Next, we compared the significantly altered proteins identified in the mouse models with data from human epilepsy samples. A total of 25 upregulated proteins were shared between the mouse and human datasets (Fig. 6A), while 14 downregulated proteins overlapped between the two species (Fig. 6B). KEGG pathway analysis revealed significant enrichment in similar pathways across both datasets, including the TCA cycle, amino acid biosynthesis, and carbon metabolism. This overlap suggests that energy metabolism dysfunction is a conserved feature of epilepsy across species, with mitochondrial disruptions potentially playing a key role in seizure activity.

**Figure 6.**
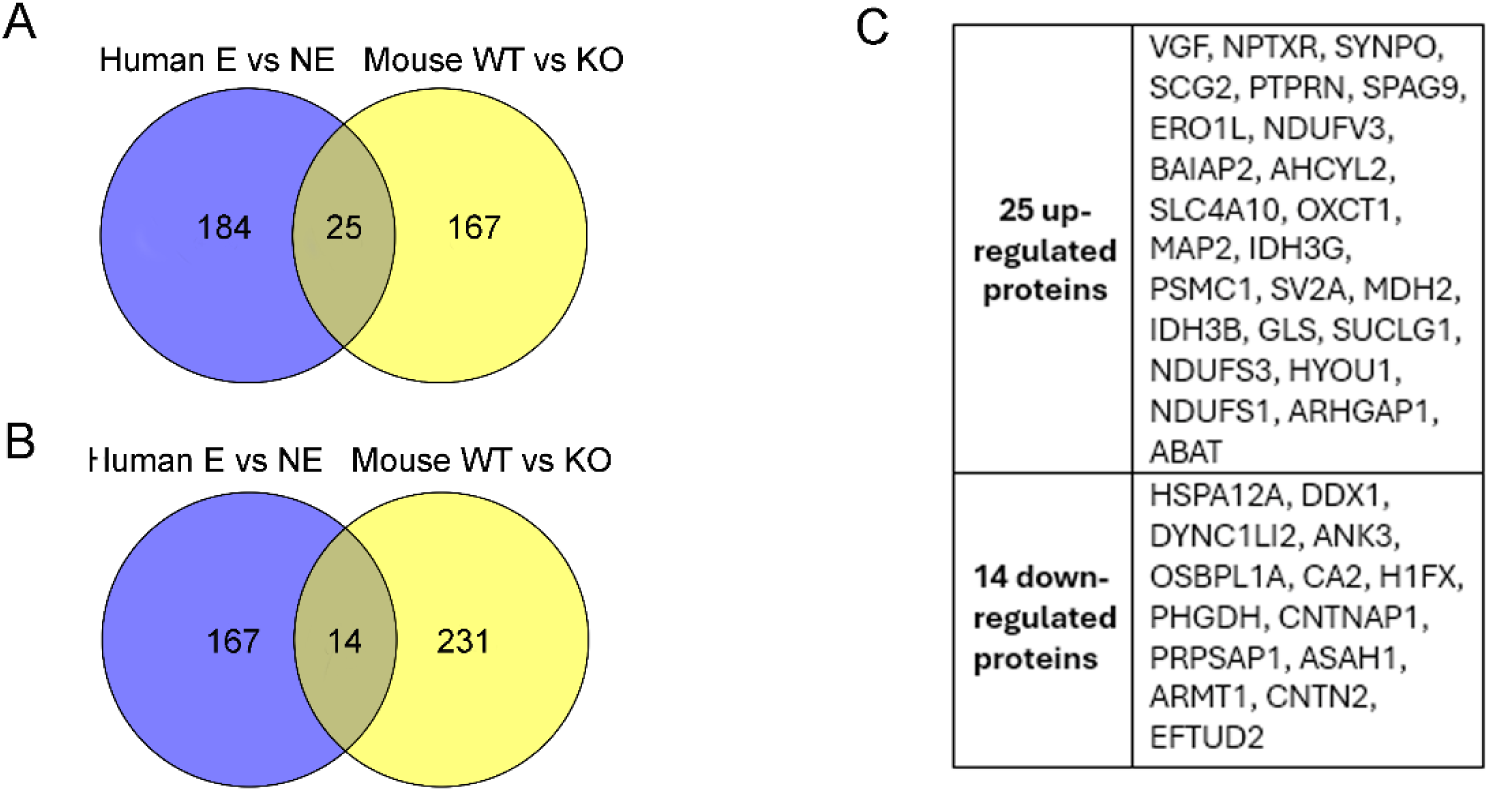
Overlapping significantly altered proteins in mouse and human epilepsy. (A) Twenty-five upregulated proteins shared between human and mouse datasets. (B) Fourteen downregulated proteins shared between human and mouse datasets. (C) Summary of overlapping proteins, including the 25 upregulated proteins shown in (A) and the 14 downregulated proteins shown in (B).

## Discussion

Using complementary pre-clinical and clinical epilepsy groups, we found significant proteomic differences highlighting both sex-dependent variations and shared molecular pathways across mouse and human epilepsy. Proteomic profiling of *Pten* KO mice and human refractory epilepsy brain samples revealed distinct clustering between epileptic and control groups (Fig. 2, Fig. 4), with prominent changes in proteins related to ion channels, metabolism, gliosis, and inflammation. The sex-specific analysis identified distinct patterns, with enrichment of protein synthesis and processing pathways in females (Fig. 3E) and enrichment of pathways associated with VEGF signaling in males (Fig. 3F), suggesting the possibility of sex-dependent epilepsy mechanisms.

Our findings align with prior omics-based epilepsy studies that have identified widespread molecular alterations between epilepsy and non-epilepsy groups in both pre-clinical models (18) and human cases (33)(for review see (16)). Previous transcriptomic and proteomic analyses consistently reported differential expression of mRNA and proteins related to synaptic function, energy metabolism, and inflammatory pathways in epileptic brain tissue compared to non-epileptic control tissues (16). We observed significant enrichment in pathways such as the TCA cycle, oxidative phosphorylation, and glutamatergic synapses in epilepsy groups (Fig. 2E), supporting a potential role for metabolic dysfunction in epilepsy. Previous studies have reported an increased accumulation of proteins linked to glutamatergic signaling in the aging mouse brain (32). The upregulation of SNCA and VGF proteins in human epilepsy samples is consistent with prior proteomic studies (33), reinforcing their potential involvement in epilepsy mechanisms.

Sex differences in seizures and epilepsy have been examined in a few omics-based studies (16), including transcriptomic analysis in hippocampi of the Scn8a N1768D genetic mouse model of pediatric epilepsy (18) and proteomic analysis of human hippocampal tissues from epilepsy cases (38). Scn8a mice develop tonic-clonic seizures by 3.5 months (18, 19), with females showing better survival, delayed seizure onset, and reduced blood-brain barrier disruption compared to males (18). At the hippocampal transcriptomic level, Scn8a females showed 15 upregulated electron transport chain genes (including Ndufb6, Ndufb7, Cox5b, and Atp5k) compared to only six downregulated genes in males (including Ndufb4, Cox7c) (18). Female *Pten* KO mice exhibited enrichment in protein processing in the endoplasmic reticulum (Fig 3E). In contrast, males in both mouse genetic epilepsy models (*Pten* and SCN8a) showed enhanced glutamatergic signaling pathways. This suggests that dysregulated glutamatergic transmission is a common feature of epilepsy across models and species, and that may be a particularly dominant pathway in males.

Detailed characterization of epilepsy progression in the *Pten* KO mouse model revealed a clear temporal pattern of seizure development. EEG analysis showed that *Pten* KO mice spent approximately 10% of their time in epileptiform activity at postnatal week 4 (25), progressively increasing to ∼40% at week 6 and ∼60% at week 9 (25), with an average mortality age of 13 weeks(25, 26). The increase in seizure activity was accompanied by progressive mossy fiber sprouting (26), enhanced astrogliosis and microgliosis, and increased mTOR pathway activation (25). While the influence of sex on seizure burden and mortality rate in *Pten* KO mice remains unknown, sex-specific differences in hippocampal microgliosis have been documented (39). At 4, 10, and 15 weeks old, male *Pten* KO mice exhibited higher microglial density within the CA1 hippocampal region compared to male WT controls, whereas female *Pten* KO mice maintained microglial density similar to female WT controls (39). These sex-specific differences align with similar male-biased inflammatory and glial responses observed in SCN8a mice (18, 19) and in models of acquired epilepsy (40) where enhanced gliosis correlated with more severe seizures in males, suggesting significant sex dimorphism in epilepsy progression.

A proteomic analysis of human epileptic hippocampal tissue surgically resected from adults with temporal lobe epilepsy (TLE) also revealed significant sex-based differences (38). Hippocampi of male patients with TLE expressed approximately 300 more specific proteins than females, showing enrichment in tumor suppressors, neurite outgrowth regulators, and Alzheimer’s disease-related proteins. In females, enrichment was documented in pathways related to synaptic plasticity, growth factors, and cytoskeletal regulators (38). While this study compared hippocampi of female and male TLE patients to sex-matched non-epileptic controls, our study compared epileptic females with epileptic males without sex-matched non-epileptic cases, which is a limitation of our study. Nevertheless, we identified 118 differentially expressed proteins enriched in metabolism-related pathways and HIF-1 signaling pathways. Although these findings further support the existence of distinct proteomes in male and female epileptic brains, it is important to note that our study examined the temporal lobe cortex rather than the hippocampus (38), so the different observations may also be attributed to the distinct tissues examined.

Metabolism-related pathways, including glycolysis, oxidative phosphorylation, and lipid metabolism, have previously been linked to epilepsy (16), with altered metabolic activity playing a crucial role in seizure generation and neuronal dysfunction (41–43). Previous studies have linked dysregulated iron metabolism to neurodegenerative conditions, including epilepsy, where excessive accumulation of iron can lead to oxidative stress and neuronal damage (44). The activation of the HIF-1 pathway, involved in iron metabolism via HO-1, may increase intracellular Fe^2+^ levels, contributing to oxidative damage and ferroptosis (45). Our study is the first to highlight the differential expression of proteins in epileptic females and males, particularly in metabolic and HIF-1 pathways.

From a mechanistic perspective, our findings reveal fundamental sex-specific responses to epileptogenic insults that extend beyond specific genetic causes. Comparative analysis across distinct mouse epilepsy models (Scn8a and *Pten* KO) and human data strongly supports convergent sex-specific pathway alterations. Female mice demonstrated enriched oxidative phosphorylation (18) and endoplasmic reticulum processing pathways (Fig. 3E), suggesting neuroprotective and/or compensatory mechanisms that may help regulate seizure severity. In contrast, males displayed a broader range of altered pathways (Fig. 3F), including those linked to glutamatergic signaling alterations likely contributing to more severe seizure phenotypes (18, 19). This mechanistic convergence indicates that sex hormones or chromosomes may fundamentally shape the brain’s response to epileptogenic triggers, directing males and females toward distinct pathophysiological processes that may result in sex-specific pathways to hyperexcitability (7, 11, 21, 22). However, our study cannot definitively determine whether these sex-specific proteomic landscapes are drivers or consequences of epileptogenesis, nor whether they serve protective or pathological roles. Taken together, our findings emphasize the critical need to analyze males and females separately in epilepsy research (7, 13), as combining sexes likely masks distinct molecular signatures that could lead to more effective sex-specific therapeutic strategies.

Several important limitations should be considered when interpreting the results of this study. While our proteomic analysis revealed distinct molecular profiles between male and female epileptic subjects, we were unable to directly correlate these profiles with seizure characteristics, as EEG monitoring was not performed in our mouse cohort. Although seizure phenotypes are well-characterized in this *Pten* KO model (25, 26), future video-EEG studies would be valuable in establishing direct relationships between sex-specific proteomic or transcriptomic signatures and differences in seizure dynamics, including both ictal and interictal activity in the same animals. The human tissue analysis also has limitations. First, our control tissues were obtained from regions adjacent to brain tumors. While these samples were histologically normal, they may have been influenced by excitatory/inhibitory imbalance in peritumoral regions (46), potentially affecting our baseline measurements. Second, the limited availability of human samples, especially non-epileptic tissues, prevented us from performing sex-matched comparisons between epileptic and non-epileptic brain biopsies. This constraint means that we cannot fully distinguish epilepsy-specific changes from baseline sex differences in the human brain biopsies. Additionally, the relatively small sample size of human brain biopsies limits the broader generalizability of our findings from the human epilepsy cases. Despite these limitations, our study provides evidence for sex-specific molecular signatures in epilepsy and highlights the critical importance of considering sex differences in both basic epilepsy research and clinical trial design.

## Author Contributions

YL: Conceptualization, Methodology, Validation, Data curation, Formal analysis, Writing – Original draft, Writing – review and editing. ZS: Writing – Original draft. KJB: Methodology, animal handling, tissue collection, genotyping. DAN: Methodology, animal handling, tissue collection, genotyping. UKA: Mass Spectrometry. JNL: Writing – review and editing. NPP: Human clinical data analysis, Writing – review, and editing. ALB: Conceptualization, Data curation, Formal analysis, Funding acquisition, Resources, Supervision, Writing – original draft, Writing – review and editing. All authors read and approved of the final manuscript.

