## Supplementary Figure 1 for "Sex-specific proteomic analysis of epileptic brain tissues from Pten knockout mice and human refractory epilepsy"

### Li et al. Supplementary Figure 1

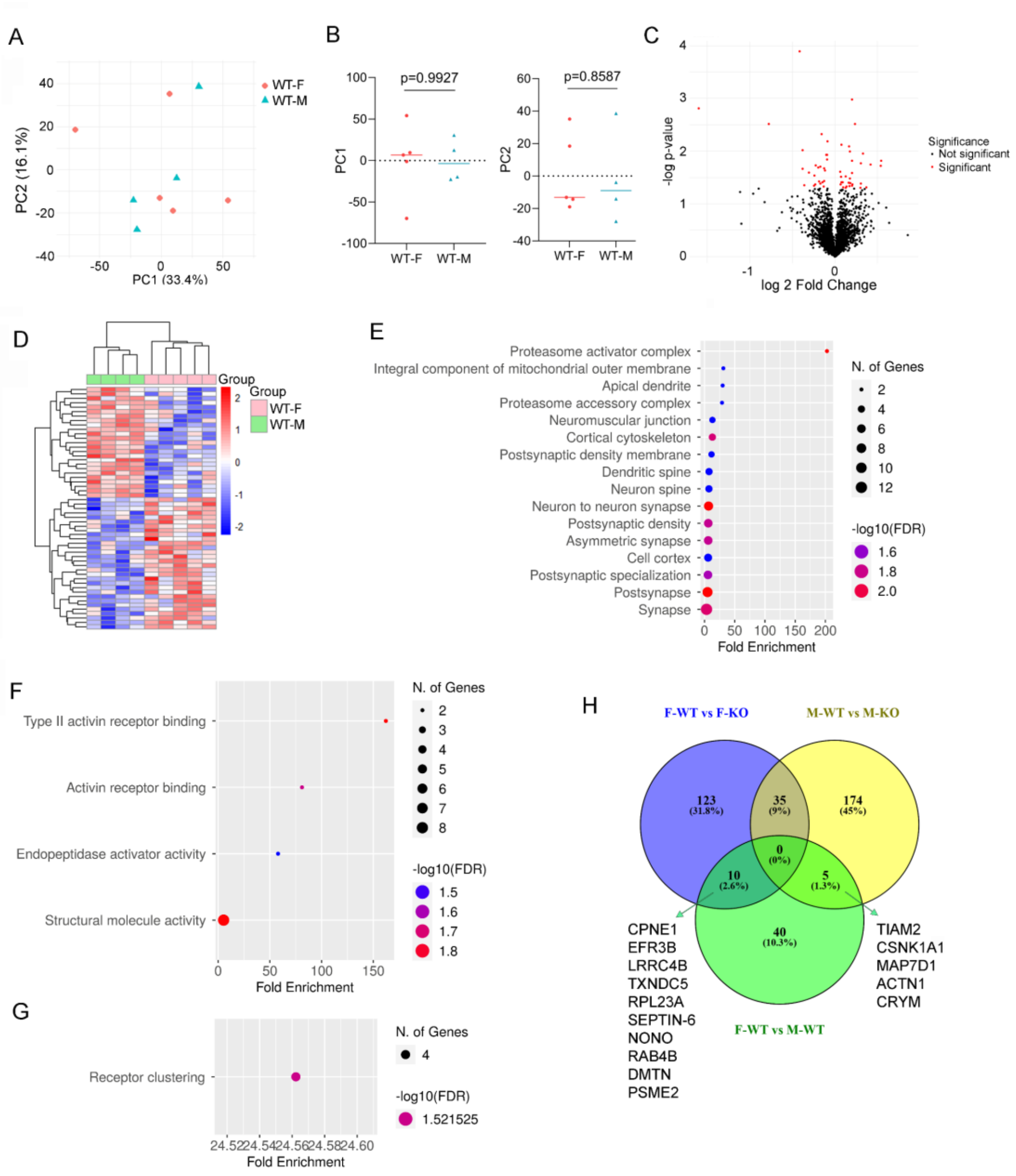


**Supplementary Figure 1. Proteomic differences between control female and male wild type (WT) mice**. (A) Principal Component Analysis (PCA) of the overall dataset. PCA of WT-F (red) and WT-M (blue) cases shows similar protein expression profiles between groups. Each point represents an individual case. (B) Bar plots of PC1 and PC2 illustrate no significant separation between WT-F and WT-M cases. (C) Volcano plot depicting protein expression differences between WT-F and WT-M mice. Proteins highlighted in red show significantly altered expression. Significance was determined using Student's t-test; corresponding p-values for significant proteins are provided in the supplementary table. (D) Heatmap of scaled protein expression values (z-scores), color-coded according to the legend, showing clustering of significantly changed proteins. (E–G) Gene Ontology (GO) analysis of significantly altered proteins, categorized by cellular component (CC) (E), molecular function (MF) (F), and biological process (BP) (G). (H) Venn diagram showing overlapping significantly altered proteins among the following comparisons: F-WT vs F-KO (blue), M-WT vs M-KO (yellow), and F-WT vs M-WT (green).
